# Host soluble inositol phosphate signaling promotes coronavirus replication

**DOI:** 10.64898/2026.09.12.751050

**Authors:** Igor Shats, Huanchen Wang, Yubai Zhou, Chunfang Gu, Adam Carr, Carlos M. Guardia, Stephen Shears, Robin Stanley, Qisheng Zhang, Raymond D. Blind, Xiaodong Wang, Xiaoling Li

## Abstract

Coronaviruses rely extensively on host pathways for replication, making host-directed therapies an attractive strategy for broad-spectrum antivirals with reduced risk of viral resistance. Here we identify the host soluble inositol phosphate pathway as a previously unrecognized dependency for coronavirus infection. Genetic or pharmacologic inhibition of several kinases in this pathway markedly suppresses replication of both alpha- and betacoronaviruses, while increasing pathway activity promotes viral replication. We developed UNC7844, a potent multi-target inhibitor of these kinases, which reduces coronavirus replication by more than four orders of magnitude in cultured cells and suppresses coronavirus infection in mice. Mechanistically, UNC7844 suppresses inositol (pyro)phosphates production, disrupts phosphoinositide homeostasis, and impairs late endosomal dynamics, blocking early post-entry steps required for viral genome release and replication. Together, our findings establish the soluble inositol (pyro)phosphate pathway as an important regulator of coronavirus infection and highlight its inhibition as a promising host-directed antiviral strategy.

## Introduction

Coronaviruses (CoVs) are enveloped, positive-sense RNA viruses that pose a persistent threat to global public health, as exemplified by the SARS-CoV, MERS-CoV, and SARS-CoV-2 outbreaks. Current antiviral strategies primarily target viral proteins; however, these approaches are often undermined by the rapid emergence of viral resistance. Targeting host pathways essential for viral replication thus offers an alternative strategy that is less susceptible to resistance and may provide broad-spectrum antiviral activity, thereby expanding the arsenal of therapeutics available for future emerging viral outbreaks and pandemics.

Soluble inositol phosphates (IPs) and their higher phosphorylated derivatives, inositol pyrophosphates, are ubiquitous signaling molecules that regulate diverse cellular processes such as vesicular trafficking, cytoskeletal dynamics, and energy homeostasis ^1, 2^. The pathway consists of multiple kinases that sequentially phosphorylate inositol 1,4,5-trisphosphate (IP3), generating higher-order inositol phosphates (IP4, IP5, and IP6), which can be further converted into inositol pyrophosphates IP7 and IP8 (Fig. 1A). Recent studies have demonstrated that IP6 is important in the life cycle of human immunodeficiency virus (HIV), where it stabilizes the viral capsid and promotes efficient viral assembly and maturation ^3–6^. These findings raise the possibility that soluble inositol phosphates may also regulate the replication of other viruses, including coronaviruses, as the entry and early replication steps of coronavirus life cycle are tightly linked to host membrane dynamics. For example, coronaviruses such as mouse hepatitis virus (MHV) and SARS-CoV-2 can enter cells via endocytic pathways, where proteolytic activation within endosomes facilitates membrane fusion and release of the viral genome into the cytosol ^7, 8^. In addition, coronaviruses extensively remodel intracellular membranes to form replication organelles enriched in specific lipid phosphoinositides, which are essential for viral RNA synthesis ^9^.

**Figure 1.**
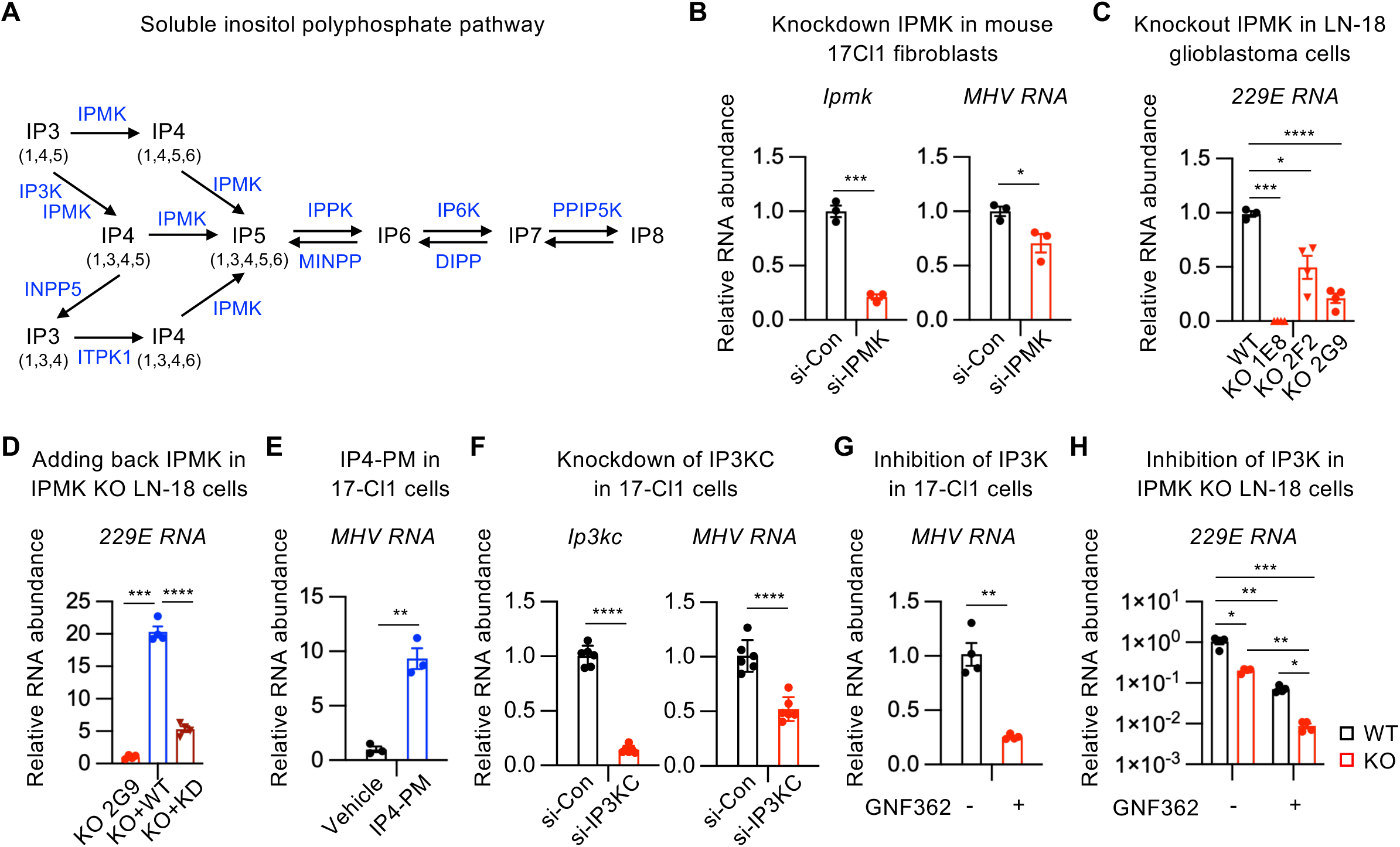
Inositol phosphate kinases promote coronavirus replication. (A) Schematic of the soluble inositol phosphate pathway. IPMK, inositol polyphosphate multikinase; IP3K, inositol-trisphosphate 3-Kinase; ITPK1, inositol-tetrakisphosphate 1-kinase; INPP5, inositol polyphosphate-5-phosphatase; IPPK, inositol-pentakisphosphate 2-kinase; IP6K, inositol hexakisphosphate kinase; PPIP5K, diphosphoinositol-pentakisphosphate kinase; DIPP, diphosphoinositol polyphosphate phosphohydrolase; MINPP, multiple inositol-polyphosphate phosphatase. (B) qPCR analysis of IPMK and MHV RNA in 17-Cl1 cells infected with MHV at an MOI of 1. Cellular RNA was collected at 24 hpi (n=3 replicates/group, values are expressed as mean ±SEM, 2-tailed unpaired t test, *p<0.05). (C) WT and IPMK KO LN-18 clones were infected with 229E virus at an MOI of 2. Virus was removed and cells were washed at 1.5 hpi. Cellular RNA was collected at 19 hpi (n=3, 4, 4, and 4 replicates, respectively, values are expressed as mean ±SEM, 2-tailed unpaired t test with Welch’s correction, *p<0.05, ***p<0.001, and ****p<0.0001). (D) Parental LN-18 IPMK KO clone 2G9 cells and its derivatives expressing WT IPMK or kinase-dead (KD) IPMK were infected with 229E at an MOI of 0.2. Virus was removed and cells were washed at 1.5 hpi. Cellular RNA was collected at 23 hpi (n=4 replicates/group, values are expressed as mean ±SEM, 2-tailed unpaired t test with Welch’s correction, ***p<0.001, and ****p<0.0001). (E) 17-Cl1 cells were infected with MHV and treated with IP4-PM or vehicle control as described in Methods. Cellular RNA was collected at 7 hpi (n=3 replicates/group, values are expressed as mean ±SEM, 2-tailed unpaired t test, **p<0.01). (F) qPCR analysis of IP3KC and MHV RNA in 17-Cl1 cells infected with MHV at an MOI of 2. Cellular RNA was collected at 16 hpi (n=6 replicates/group, values are expressed as mean ±SEM, 2-tailed unpaired t test, ****p<0.0001). (G) 17-Cl1 cells were infected with MHV at an MOI of 6. Viruses were removed, cells were washed and treated with 5 μM GNF362 or vehicle control at 1.5 hpi. Cellular RNA was collected at 24 hpi (n=4 replicates/group, values are expressed as mean ±SEM, 2-tailed unpaired t test, **p<0.01). (H) WT LN-18 cells and IPMK KO clone 2G9 were infected with 229E at an MOI of 1. Viruses were removed, cells were washed and treated with 5 μM GNF362 or vehicle control at 1.5 hpi. Cellular RNA was collected at 24 hpi (n=4 replicates/group, values are expressed as mean ±SEM, 2-way ANOVA, *p<0.05, **p<0.01, and ***p<0.001).

In the present study, we demonstrate that two key kinases in the soluble inositol (pyro)phosphate pathway, inositol polyphosphate multikinase (IPMK) and inositol trisphosphate kinase (IP3K), promote coronavirus replication. UNC7844, a novel multi-target inositol phosphate kinase inhibitor, potently suppresses replication of coronaviruses in vitro and in vivo. Mechanistically, UNC7844 inhibits early stages of the viral life cycle by disrupting viral trafficking and phospholipid remodeling. Our findings identify the soluble inositol phosphate pathway as a promising new host-directed anti-coronavirus target and disclose efficient lead compounds for therapeutic development.

## Results

### Inositol phosphate kinases promote coronavirus replication

To examine the potential role of the host soluble inositol (pyro)phosphate pathway in the life cycle of coronaviruses, we first investigated host IPMK, an enzyme that catalyzes multiple steps in the conversion of IP3 to IP4 and IP5 (Fig. 1A). In mouse 17-Cl1 fibroblasts, knockdown of IPMK significantly inhibited the replication of mouse hepatitis virus (MHV), a coronavirus that belongs to the same betacoronavirus family as SARS-CoV-2 (Fig. 1B). Consistently, knockout of IPMK in human LN-18 glioblastoma cells significantly inhibited the replication of human conventional alphacoronavirus 229E in three independent knockout clones (Fig. S1A-S1C, and 1C). Conversely, overexpression of WT IPMK in an IPMK KO clone dramatically enhanced viral replication (Fig. S1D and 1D, WT IPMK), whereas overexpression of the kinase-dead D144A IPMK mutant produced a significantly weaker effect (Fig. S1D and 1D, KD IPMK). Moreover, IP4-PM, a cell-permeable form of IP4 which is hydrolyzed by the cellular esterases to release free IP4(1,3,4,5) upon cell entry, strongly enhanced MHV replication in MHV-infected 17-Cl1 cells (Fig. 1E). These observations demonstrate a proviral role of IPMK in both human and mouse cells.

We then evaluated the role of IP3K activity in coronavirus replication. Knockdown of IP3KC in 17-Cl1 cells significantly inhibited the replication of MHV (Fig. 1F). Furthermore, pharmacologic inhibition of IP3K activity with GNF362 significantly decreased MHV RNA in MHV-infected 17-Cl1 cells (Fig. 1G). GNF362 also suppressed the replication of 229E by 10-fold in WT LN-18 cells (Fig. 1H, WT+GNF362 vs WT-GNF362). Notably, inhibition of IP3K in IPMK KO LN-18 cells led to a more than 100-fold decrease in viral RNA compared with parental cells (Fig.1H, KO+GNF362 vs WT-GNF362). Taken together, these results indicate that the soluble inositol phosphate pathway promotes coronavirus replication and further suggest that simultaneous inhibition of several kinases in this pathway could highly efficiently suppress coronavirus.

### UNC7844 is a multi-target inositol phosphate kinase inhibitor

To test whether pharmacological inhibition of the soluble inositol (pyro)phosphate metabolism could suppress coronavirus replication, we developed UNC7844, a new small molecule inhibitor based on two recently reported parental compounds, UNC7437 and UNC7467. UNC7437 was first reported as an IPMK inhibitor, displaying IC₅₀ values of 12.8 nM for IPMK, 146 nM for IP6K2, and 8 µM for IP3KA (Fig. 2A) ^10, 11^. UNC7467 exhibited approximately a 10-fold higher potency against IP3KA and a 30-fold higher potency against IP6K2 relative to UNC7437 (Fig. 2B) ^12^. Further refinement yielded UNC7844 that displayed additional 26-fold increase in potency for IP3KA, a 5-fold improvement for IPMK, and a 3-fold increase for IP6K2 (Fig. 2C and 2D). These optimizations positioned UNC7844 as a highly potent inhibitor of several kinases in the soluble inositol (pyro)phosphate pathway.

**Figure 2.**
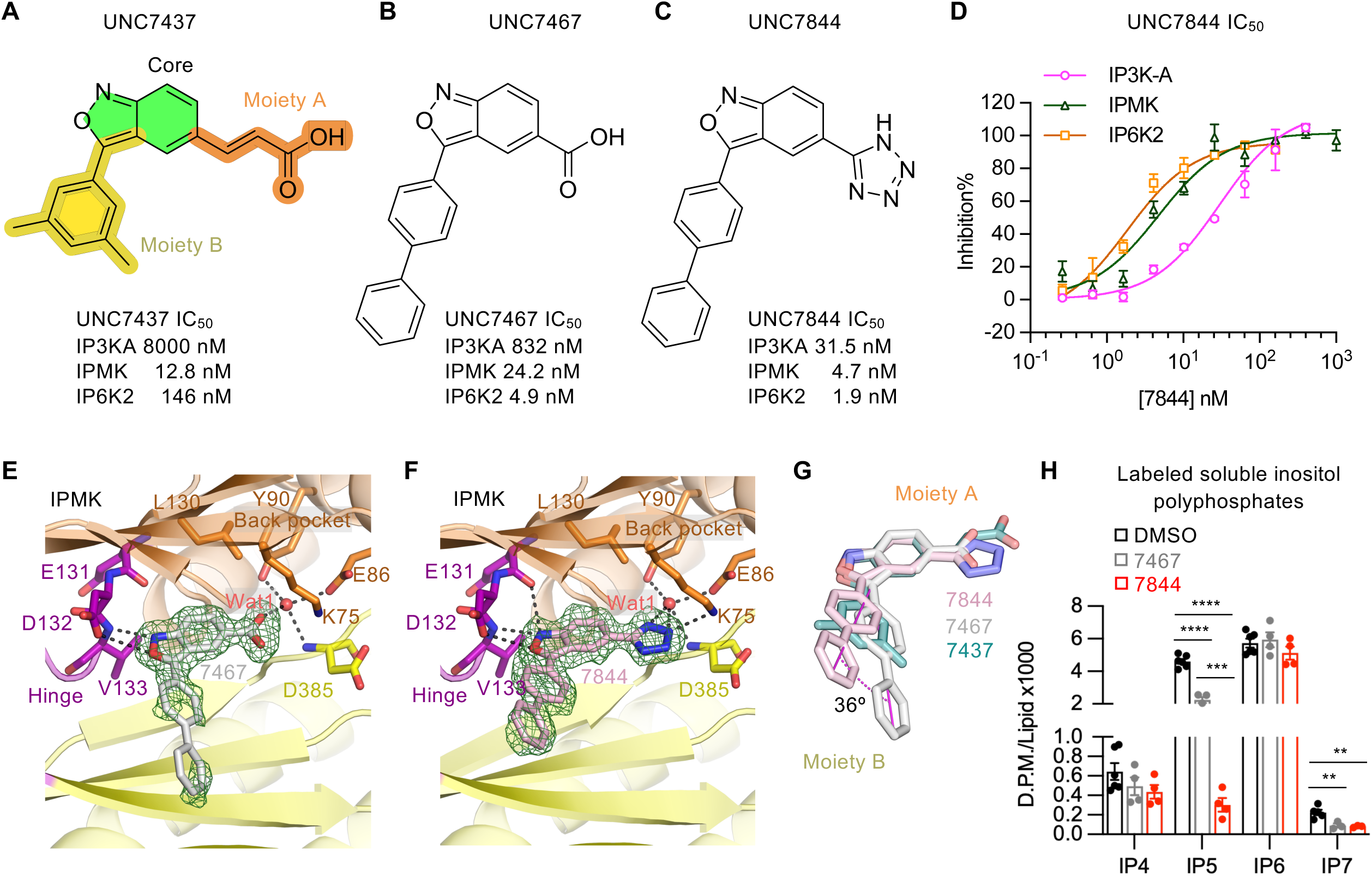
UNC7844 is a potent multitarget inhibitor of soluble inositol phosphate kinases. (A–C) Chemical structures and IC₅₀ values of UNC7437, UNC7467, and UNC7844. (D) In vitro kinase assays used for determination of IC₅₀ values of UNC7844 against IP3K-A, IPMK, and IP6K2 (n=3 replicates/group, values are expressed as mean±SEM). (E–F) Structural basis for the binding of UNC7467 and UNC7844 to IPMK as determined by X-ray crystallography. The difference Fo–Fc electron density maps are contoured at 2.5 σ and shown in green mesh. (G) Structural overlay of UNC7437, UNC7467, and UNC7844 bound to IPMK. (H) Quantification of intracellular inositol phosphate levels following overnight treatment (10 µM) with UNC7467 or UNC7844 (n=6 replicates DMSO, 4 replicates UNC7644, and 4 replicates UNC7844, values are presented as mean±SEM, 2-tailed unpaired t test with Welch’s correction, **p<0.01, ***p<0.001, and ****p<0.0001).

Analysis of crystal structures of UNC7467 (Fig. 2E) and UNC7844 (Fig. 2F) in complex with the IPMK catalytic core domain (Supplementary Table 1), alongside the previously reported UNC7437–IPMK complex ^10^, revealed that all three inhibitors occupy the ATP-binding site of IPMK, forming canonical hydrogen bonds with the hinge region while exploiting the “KEY” motif in the back pocket to enhance potency through additional polar contacts with protein residues and structured water molecules (Fig. 2E and 2F). Notably, UNC7844 formed one additional hydrogen bond within the hinge and two additional hydrogen bonds with the KEY motif compared with UNC7467, accounting for its increased potency against IPMK. This rearrangement also induced a ∼36° rotation of moiety B (Fig. 2G), which might explain the dramatic improvement in IP3K-A inhibition and enhanced activity against IP6Ks.

Since UNC7844 occupies the ATP binding pocket of the inositol kinases, its selectivity was evaluated in a panel of 30 kinases that provided a broad survey of protein kinase families. Only seven kinases were inhibited by more than 50% at 1 µM UNC7844 (Supplementary Table 2), demonstrating that UNC7844 exhibits significant, albeit incomplete, selectivity towards soluble inositol phosphate kinases, with IC_50_ in the low nanomolar range (Fig. 2C and 2D).

Further cellular metabolic labeling experiments showed that both UNC7467 and UNC7844 moderately decreased the IP4 levels and highly significantly reduced IP5 and IP7 (Fig. 2H). However, UNC7844 produced a much greater decrease in IP5 levels than UNC7467 (Fig. 2H), demonstrating that UNC7844 is a more potent inhibitor of soluble inositol phosphate kinases compared to the parental compound in cellular assays. Notably, cellular levels of IP6 remained unchanged under these conditions, suggesting that the cellular content of IP6 is relatively stable and less susceptible to acute perturbation by these inhibitors.

### Inositol phosphate kinase inhibitors suppress coronavirus infection

Next, we examined the effects of these inhibitors on coronavirus replication. UNC7467 dose-dependently inhibited 229E replication in THP1-derived human macrophages, with an IC_50_ of approximately 4 μM (Fig. S2A). Notably, preincubation with 10 μM UNC7467 for 3 days followed by compound washout prior to 229E infection did not affect viral replication (Fig. S2B), suggesting that UNC7467 does not act by activating intrinsic cellular antiviral mechanisms. Instead, UNC7467 must be present during infection to exert its antiviral effects. Interestingly, we observed that increasing medium fetal bovine serum (FBS) concentration to 10% strongly decreased the antiviral activity of all tested compounds, whereas complete removal of FBS enhanced their antiviral potency. Consequently, all subsequent cell culture experiments in this study were performed in FBS-free medium.

As shown in Fig. 3A and S2C, UNC7437 and UNC7467 reduced 229E RNA by 97% at 10 μM and by more than 100-fold at higher concentrations in THP1-derived macrophages (TDMs). Remarkably, UNC7844, the multi-kinase inhibitor that potently inhibits IPMK, IP6K and IP3K (Fig. 2C), was substantially more potent, decreasing 229E RNA by 99.9% at 10 μM (Fig. 3A and S2C). None of the inhibitors significantly reduced cell viability of TDMs (Fig. S2D). In the MHV infection model in mouse 17-Cl1 fibroblasts, UNC7437 and UNC7467 decreased viral RNA by 55% and 60%, respectively, after 4 hours of treatment at 10 μM (Fig. 3B). UNC7844, again, was the most efficient compound among the three, resulting in a four-order-of-magnitude reduction of MHV RNA at 10 μM in 17-Cl1 fibroblasts (Fig. 3B) and 93% inhibition of MHV replication in primary mouse bone marrow-derived macrophages (BMDMs), where UNC7467 was inactive at 10 μM (Fig. 3C). None of the inhibitors displayed strong cytotoxicity in uninfected 17-Cl1 cells (Fig, 3D, No virus), and importantly, UNC7437 and UNC7844 completely rescued MHV-induced cell death at 10 μM (Fig. 3D, MHV vs no virus).

**Figure 3.**
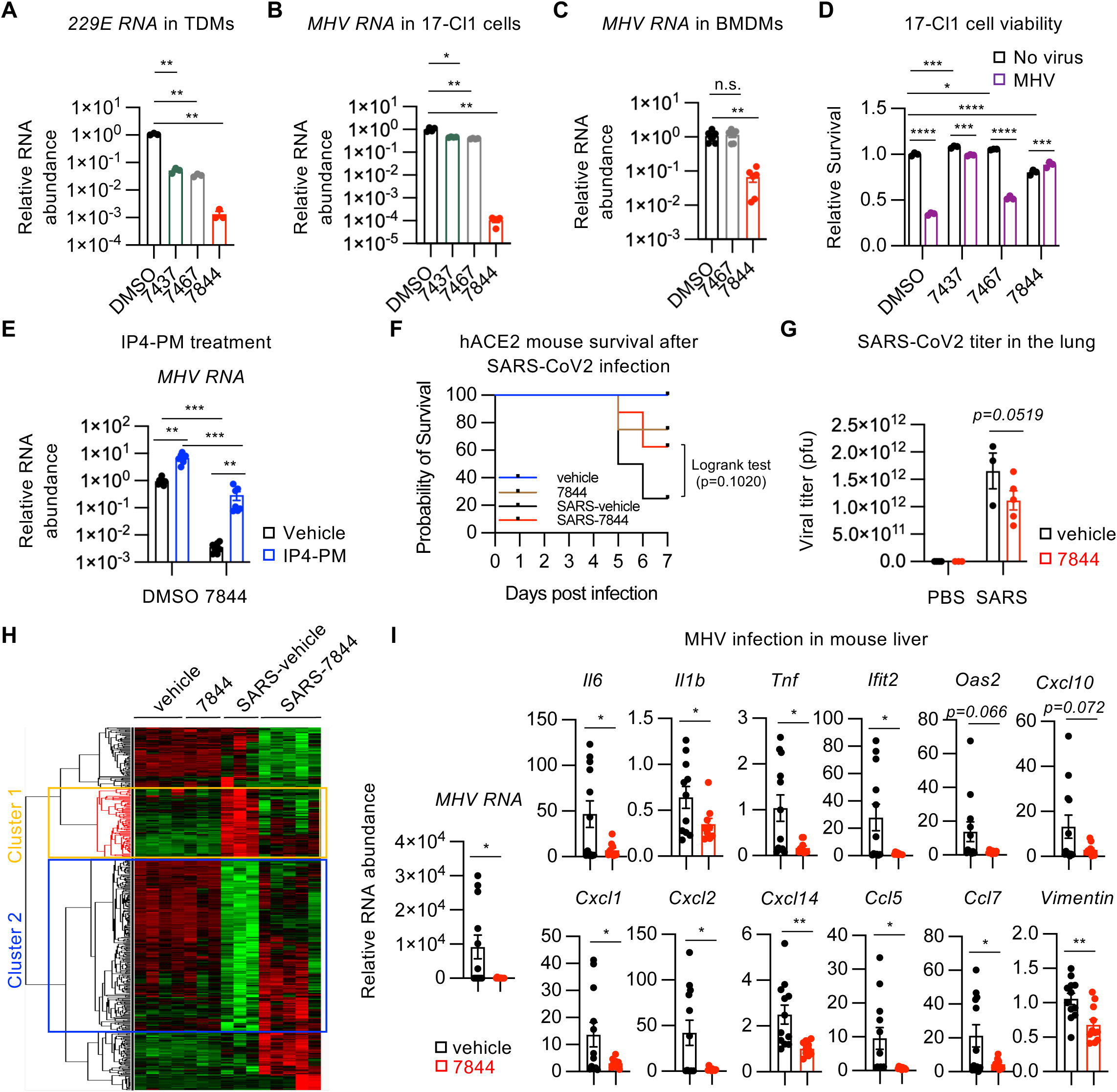
Pharmacologic targeting of inositol phosphate kinases inhibits coronavirus replication in vitro and in vivo. (A) THP1 cells were differentiated with 50 nM phorbol myristate acetate (PMA) for 24 hours, and the resulting macrophages (TDMs) were infected with 229E at an MOI of 1. Virus was removed at 2 hpi and cells were washed and treated with the indicated inhibitors at 10 μM for 17 hours (n=3 replicates/group, values are presented as mean±SEM, 2-tailed unpaired t test with Welch’s correction, **p<0.01). (B) 17-Cl1 cells were infected with MHV at an MOI of 0.2 and virus was removed at 2 hpi. Cells were washed and treated with the indicated inhibitors at 10 μM, then harvested at 6 hpi (n=4 replicates/group, values are presented as mean±SEM, 2-tailed unpaired t test with Welch’s correction, *p<0.05, **p<0.01). (C) BMDMs were infected with MHV-S at an MOI of 0.5. Cellular RNA was collected at 21 hpi (n=6 replicates/group, values are expressed as mean ±SEM, 2-tailed unpaired t test with Welch’s correction, n.s., not significant, **p<0.01). (D) Cell viability of 17-Cl1 cells treated as in (B) was determined by CellTiter-Glo assay at 44 hpi (n=3 replicates/group, values are expressed as mean ±SEM, 2-way ANOVA, *p<0.05, ***p<0.001, ****p<0.0001). (E) 17-Cl1 cells were infected with MHV-GFP virus at an MOI of 2. Virus was removed at 1 hpi and cells were washed and treated with 10 μM cell-permeable form of IP4 (IP4-PM), 10 μM UNC7844 or their combination as described in Methods. Cellular RNA was collected at 7 hpi. Results are from two independent experiments each performed in biological triplicates (n=6 replicates/group, values are presented as mean±SEM, 2-Way ANOVA, **p<0.01, ***p<0.001). (F-H) UNC7844 shows trend for antiviral activity against SARS-CoV-2 *in vivo*. hACE2-overexpressing mice were intranasally infected with SARS-CoV-2 and treated with 10 mg/kg twice daily UNC7844 or vehicle control for 7 days (Animal numbers: initial 4, 4, 8, and 8 mice; final 4, 3, 3, and 5 mice, respectively). (F) Survival curves of experimental mice, logrank test. (G) Viral titers in the lung were determined by plaque assay (n= 4, 3, 3, and 5 mice, respectively; values are expressed as mean ±SEM, 2-tailed unpaired t test between 7844 vs vehicle treated SARS infected mice). (H) Heatmap of genes differentially expressed in the lungs of UNC7844- and vehicle-treated infected mice. (I) UNC7844 inhibits MHV replication and suppresses inflammation *in vivo*. Mice were infected with 10^6^ pfu MHV i.p. and treated with 25 mg/kg i.p. UNC7844 or vehicle control twice daily for 48 hours. (n=12 mice in vehicle group and n=10 mice in UNC7844 group, values are presented as mean±SEM, 2-tailed unpaired t test, *p<0.05, **p<0.01).

We then tested whether the antiviral activity of UNC7844 is mediated through inhibition of the soluble inositol (pyro)phosphate pathway. Consistent with the observed proviral activity of this pathway (Fig. 1), IP4-PM elevated MHV RNA abundance in MHV-infected 17-Cl1 cells (Fig. 3E, DMSO). Importantly, IP4-PM partially reversed the antiviral activity of UNC7844 (Fig. 3E, 7844), suggesting that inhibition of IPMK/IP3K-mediated IP4 synthesis significantly contributes to the antiviral effect of the inhibitor.

UNC7844 also displayed strong antiviral activity in vivo. Following intravenous administration (3 mg/kg), UNC7844 had a t_1/2_ of 0.6 hours in mice (Supplementary Table 3), suggesting a need for several doses per day. In a pilot study using SARS-CoV-2-infected hACE2-overexpressing mice, twice daily treatment with 10 mg/kg UNC7844 for 7 days showed a trend toward reduced mortality. Five out of eight mice survived in the UNC7844-treated group compared with three of eight mice in the vehicle-treated group (Fig. 3F, log-rank test p=0.1). UNC7844-treated mice also showed reduced SARS-CoV-2 titer and mRNA levels of inflammatory markers *Il6* and *Tnf* in the lung, although these results did not reach statistical significance (Fig. 3G and S2E). Further genome-wide transcriptomic analysis of lung tissues revealed that 7-day SARS infection induced a cluster of genes involved in cell cycle process and pre-mRNA splicing (Fig. 3H and S2F, Cluster 1), while reducing the expression of a separate cluster of genes associated with microtubule-based transport and cilium movement (Fig. 3H and S2F, Cluster 2). UNC7844 treatment attenuated the SARS-induced changes in both gene clusters (Fig. 3H), consistent with suppression of viral activity. These promising results prompted us to perform dose escalation experiments, which suggested that UNC7844 dose can be increased to 25 mg/kg twice daily without significant body weight loss (Fig. S2G). At this dose, UNC7844 significantly reduced MHV RNA levels and suppressed the expression of multiple cytokines and chemokines as well as fibrosis marker *vimentin* in the liver of MHV A59-infected C57BL/6J mice (Fig. 3I). Together, our findings demonstrate that pharmacologic targeting of the soluble inositol phosphate pathway inhibits coronavirus replication in vitro and in vivo and further suggest that simultaneous inhibition of multiple kinases in this pathway, as exemplified by UNC7844, yields a more robust antiviral effect.

### UNC7844 targets endocytic viral trafficking during the early stages of viral infection

To define how inhibition of the soluble inositol phosphate pathway suppresses coronavirus replication, we analyzed the kinetics of MHV infection in 17-Cl1 cells. In control MHV-infected 17-Cl1 cells, viral RNA rose exponentially following removal of the extracellular virus at 2 hours post-infection (hpi), with 4000-fold increase by 7 hpi (Fig. 4A, DMSO). Addition of UNC7844 at 2 hpi completely prevented this rise (Fig. 4A, 7844). At this time point, coronavirus internalization via endocytosis is largely complete, although endosomal trafficking and membrane fusion may still be ongoing ^7, 8^. Therefore, UNC7844 likely inhibits early stages of infection after viral entry.

**Figure 4.**
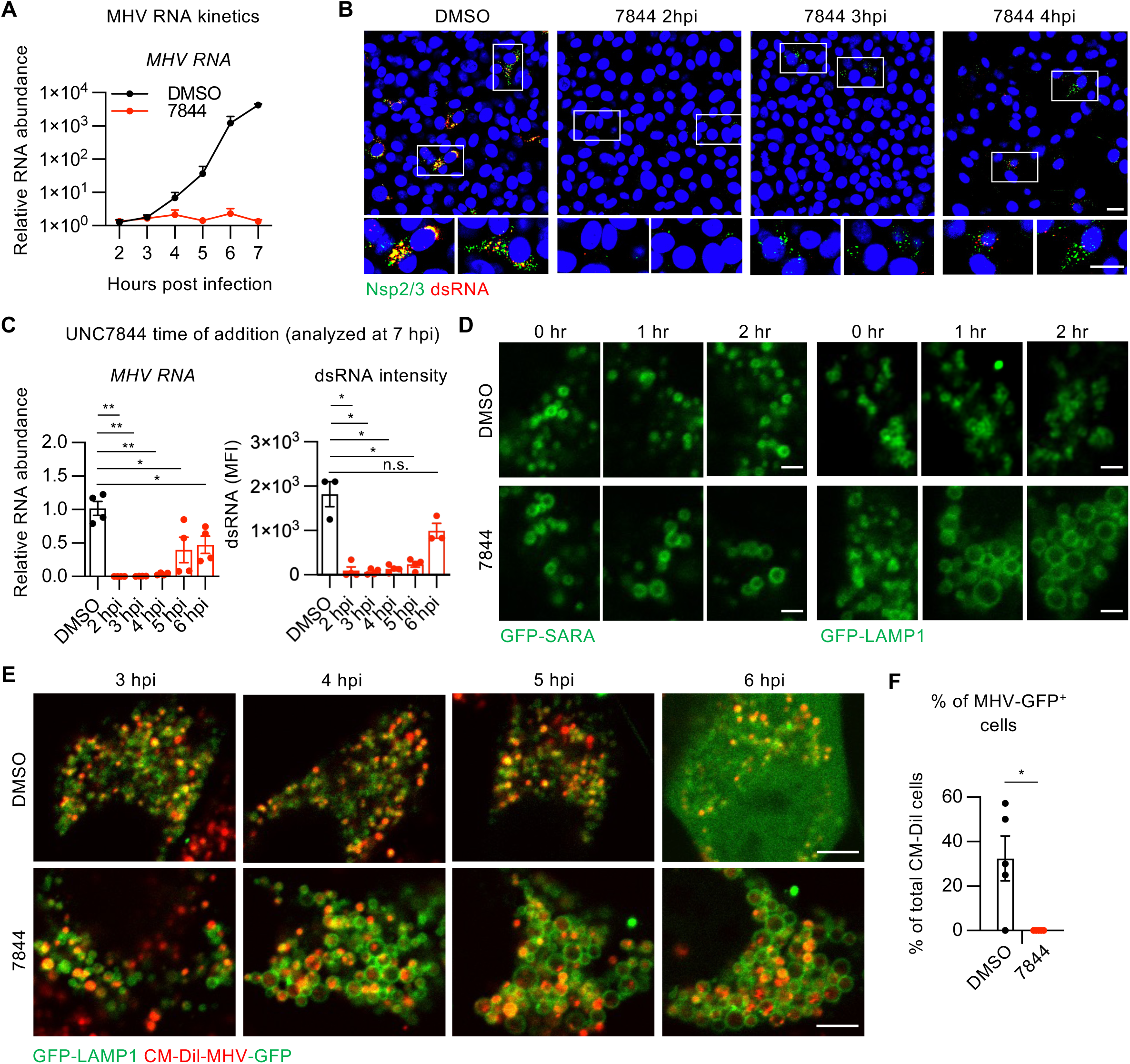
UNC7844 targets early events in coronavirus life cycle. (A) 17-Cl1 cells were infected with MHV at an MOI of 7. Virus was removed at 1 hpi and cells were washed and treated with 10 μM UNC7844 or DMSO control. Cellular RNA was collected at indicated time points (n=2 replicates/group, values are presented as mean±SEM). (B) 17-Cl1 cells were infected with MHV at an MOI of 5 for 1 hour, washed and treated with 10 μM UNC7844 at indicated time points. Cells were fixed at 5 hpi and stained for nsp2/3 (green), dsRNA (red). Scale bars, 20 μm. (C) Experiment was performed similarly to (B) but RNA was collected (left, n=4 replicates/group) or cells were fixed and stained at 7 hpi and intensity of dsRNA signal quantified (right, n=3 replicates/group). Values are presented as mean± SEM, 2-tailed unpaired t test with Welch’s correction, *p<0.05, **p<0.01. (D) 17-Cl1 cells were transiently transfected with GFP-SARA or GFP-LAMP1, treated with 10 μM UNC7844 and monitored by time-lapse microscopy. Scale bars, 2 μm. (E) Experiment was performed like in (D) but cells were infected with MHV-GFP virions labeled with CM-DiI dye (red) and treated with 10 μM UNC7844 at 3 hpi. The diffuse green fluorescence at 6 hpi in DMSO-treated cells reflects the expression of the virally encoded GFP. Scale bars, 5 μm. (F) Quantification of the percentage of CM-DiI-positive cells that expressed GFP at 6 hpi (n=5 independent fields/group, values are presented as mean± SEM, 2-tailed unpaired t test, *p< 0.05).

To further characterize these early stages in the virus life cycle, we performed time of addition experiments where we infected the cells with MHV for one hour, washed the virus, then added UNC7844 at 2, 3, or 4 hpi before fixing the cells at 5 hpi. Immunofluorescence staining for a viral non-structural protein nsp 2/3 (marker for viral protein translation) and for dsRNA (marker of active viral replication) revealed that a significant fraction of DMSO-treated cells had multiple nsp2/3 (green), dsRNA (red) or double-positive (yellow) foci corresponding to active replication organelles at 5 hpi (Fig. 4B, DMSO). In contrast, in cells treated with UNC7844 at 2 hpi, both nsp2/3 staining and dsRNA RNA-positive foci were almost completely absent at 5 hpi (Fig. 4B, 7844 2hpi), suggesting an almost total block of viral protein translation and complete block of viral replication. When added at 3 or 4 hpi, UNC7844 still completely blocked the appearance of dsRNA but nsp2/3 expression was progressively less affected (Fig. 4B, 7844 3hpi, 4hpi). In a separate experiment, addition of UNC7844 as late as 5 hpi still significantly reduced viral RNA levels and dsRNA staining intensity at 7 hpi (Fig. 4C). These findings suggest that UNC7844 may affect several stages of the viral life cycle, potentially including endosomal trafficking as well as post-uncoating processes such as formation or function of the replication organelles.

To identify the affected trafficking step, we monitored vesicle dynamics by live-cell imaging in 17-Cl1 cells. UNC7844 at 10 μM selectively enlarged LAMP1-positive late endosomes/lysosomes without markedly affecting SARA-positive early endosomes (Fig. 4D). Time-lapse analysis of these cells infected with CM-Dil-labeled MHV-GFP virions further revealed that by 3 hpi, viral particles were largely outside early endosomes and localized to late endo/lysosomes (Fig. S3A, S3B, and 4E, 3hpi). In DMSO-treated cells, LAMP1-positive endo/lysosomes exhibited rapid dynamics, accompanied by diffuse expression of the virally encoded GFP in a substantial fraction of infected cells by 6 hpi (Fig. 4E, 4F, S3B, and Supplementary Movie 1), indicating successful virion uncoating and access of the viral genomic RNA to the host translation machinery. In contrast, following UNC7844 treatment at 3 hpi, the enlarged late endosomes/lysosomes displayed markedly slower dynamics, and virions remained trapped within these vesicles, failing to reach the cytosol and thereby preventing translation of viral-encoded GFP (Fig. 4E, 4F, S3B, and Supplementary Movie 2). Collectively, these results are consistent with the notion that UNC7844 inhibits endocytic viral trafficking/uncoating.

### Modest lipid inositol kinase inhibition and phosphoinositide disruption by UNC7844 does not fully account for its antiviral activity

The ability of UNC7844 to inhibit viral RNA synthesis even when added at 4-5 hpi, after the expression of viral proteins and the replication of vial RNA (Fig. 4B and 4C), suggested that it disrupts post-entry stages beyong endocytic trafficking, potentially including replication organelle formation. Since these processes involve extensive lipid remodeling, we compared lipidomes of control and UNC7844-treated 17-Cl1 cells (Supplementary Table 4). The top upregulated lipids in UNC7844 treated cells were highly enriched for multiple phosphatidylinositols (PI) (Fig. 5A and 5B). Because our LC-MS method measured the abundance of the unphosphorylated species, these results suggest that UNC7844 may disrupt the equilibrium among different phosphoinositide species, leading to accumulation of the unphosphorylated PIs. Indeed, immunofluorescence staining for different phosphoinositides with specific antibodies confirmed that UNC7844 decreased the staining intensity of PI(3)P, PI(4)P and PI(4,5)P_2_ (Fig. 5C and S4A). In addition, in DMSO-treated cells, the staining signal of PI(3,4,5)P₃ was predominantly localized to cytosolic puncta, likely in endosomal compartments. In UNC7844 treated cells, in contrast, PI(3,4,5)P₃ staining was redistributed to the plasma membrane (Fig. 5C and S4A), suggesting disruption of endosomal trafficking and phosphoinositide compartmentalization.

**Figure 5.**
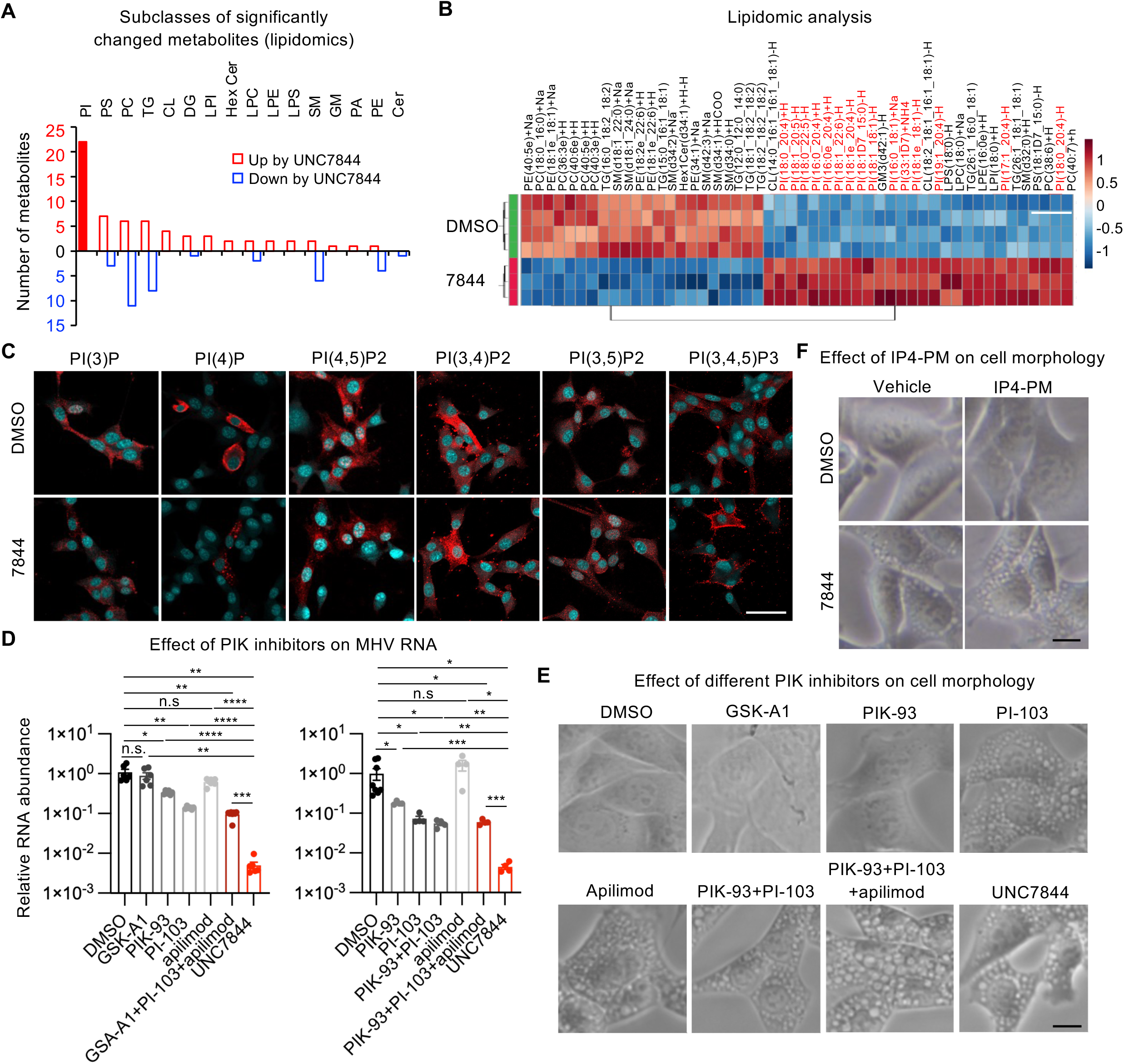
UNC7844 remodels phosphatidylinositide composition and exhibits substantially greater antiviral efficacy than selective phosphoinositide kinase inhibitors. (A and B) UNC7844 remodels phosphatidylinositide composition. 17-Cl1 cells were treated with 10 μM UNC7844 or DMSO control for 4 hours and analyzed by LC-MS-based lipidomics. (A) Differential lipids were categorized by class and the number of up- and downregulated specific lipids for each lipid class is plotted. (B) Heatmap of the top 50 differential lipids. PIs are highlighted in red. (C) Immunostaining for specific phosphoinositides. Bar, 50 μm. (D) Side-by-side comparison of antiviral activities of different PIK inhibitors with UNC7844. 17-Cl1 cells were infected with MHV at an MOI of 3. Virus was removed at 1 hpi, cells were washed and treated with 10 μM GSK-A1, 10 μM PIK-93, 10 μM PI-103, 200 nM apilimod, indicated combinations, or 10 μM UNC7844 for 4 hours (n=4-6 replicates/group, values are presented as mean±SEM, 2-tailed unpaired t test with Welch’s correction, *p<0.05, **p<0.01, ***p<0.001, ****p<0.0001, n.s., not significant). (E) Effect of different PIK inhibitors and UNC7844 on the morphology of 17-Cl1 cells. Cells were treated as in (D) and analyzed at 5 hpi. Bar, 10 μm. (F) Effect of IP4-PM on the morphology of 17-Cl1 cells. 17-Cl1 cells were infected with MHV-GFP virus and treated with 10 μM IP4-PM, 10 μM UNC7844 or their combination as described in Methods. Cells were analyzed at 6.5 hpi. Bar, 10 μm.

Phosphoinositide kinases and phosphatases tightly regulate phosphoinositide homeostasis ^13^, and the kinases in particular have been implicated in replication of positive-strand RNA viruses including coronaviruses ^14–18^. The observed changes in phosphoinositide metabolism raised the possibility that UNC7844 may suppress coronavirus replication partially through direct inhibition of phosphoinositide kinases. To test this possibility, we profiled UNC7844 against a panel of 26 lipid inositol kinases. At 1 μM, UNC7844 inhibited PI4KA, PI4KB, PIKfyve, and PI3KC2G by more than 50% (Supplementary Table 5), although it remained substantially more potent toward soluble inositol phosphate kinases (Fig. 2C and 2D). We then directly compared the antiviral effects of UNC7844 with selective inhibitors of PI4KA (GSK-A1), PI4KB (PIK-93), PI3K (PI-103), and PIKfyve (apilimod), individually and in combination, each at 10 μM except apilimod, which was used at 200 nM. None of these treatments reproduced the antiviral potency of UNC7844 (Fig. 5D).

We further tested the possible interactions between UNC7844 and lipid kinase inhibitors described above, as well as additional selective inhibitors of lipid-embedded phosphoinositide kinases using lower concentrations commonly employed to achieve efficient yet target-specific inhibition. UNC7844 was used at 5 μM, which resulted in only partial antiviral effect, thereby providing a window for detecting further modulation by the lipid kinase inhibitors. At these lower concentrations, with the exception of GDC980, a class I PI3K inhibitor that only enhanced viral replication both alone or in combinations with UNC7844, the other specific lipid kinase inhibitors had minimal impact on viral replication when used alone and did not significantly alter the antiviral activity of UNC7844 (Fig. S4B-S4D).

The PIKfyve-specific inhibitor apilimod has previously been shown to disrupt endosomal trafficking and to have an antiviral activity against various viruses, including filoviruses and coronaviruses ^17, 19^. We found that UNC7844, PI-103, and inhibitor combinations also induced extensive cytosolic vacuoles resembling those observed with apilimod (Fig. 5E). However, the antiviral activity of PI-103, apilimod, and the inhibitor combinations remained substantially weaker than that of UNC7844 (Fig. 5D). Moreover, the partial reversal of the antiviral activity of UNC7844 by IP4-PM (Fig. 3E) was not associated with any reduction in cytosolic vacuolization (Fig. 5F). Thus, UNC7844-induced endolysosomal vacuolization and any potential inhibition of lipid inositol kinases PI3K, PI4K, or PIKfyve are unlikely to be solely responsible for its antiviral activity. Together with the strong proviral activity of IP4-PM, these findings support inhibition of the soluble inositol phosphate pathway as the predominant mechanism underlying the antiviral activity of UNC7844.

## Discussion

Our study identifies the soluble inositol phosphate pathway as a previously unrecognized host pathway required for efficient coronavirus replication. Genetic and pharmacological inhibition of inositol phosphate kinases consistently suppressed viral replication, whereas enhancing the activity of this pathway increased infection, demonstrating direct proviral role of soluble inositol phosphates. Importantly, simultaneous inhibition of multiple kinases by UNC7844 produced substantially greater antiviral effect than targeting individual enzymes, suggesting that coronavirus replication depends on the integrated output of the soluble inositol phosphate network rather than on any single kinase.

Our data further indicate that UNC7844 acts during the early stages of coronavirus infection. Time-of-addition experiments, together with live-cell imaging, demonstrate that UNC7844 disrupts late endosomal trafficking, preventing efficient release of the viral genome into the cytosol (Fig. 4). These observations, combined with the central role of endocytosis in viral entry and maturation as well as the established function of inositol pyrophosphates in regulating endocytosis ^20^, supports a model in which inhibition of endocytic pathways contributes to the antiviral activity of UNC7844. This is consistent with previous studies showing that coronaviruses rely on endosomal maturation and protease activity to complete membrane fusion and release their genome (Burkard et al., 2014; Bayati et al., 2021). However, several observations indicate that impaired trafficking alone cannot fully account for the remarkable antiviral efficacy of UNC7844. First, selective inhibition of PI3K, PI4K, or PIKfyve induced similar vacuolar phenotypes but failed to reproduce the profound inhibition of coronavirus replication achieved by UNC7844 (Fig. 5D and 5E). Secondly, UNC7844 retained substantial antiviral activity even when added after viral uncoating had largely occurred (Fig. 4B and 4C), suggesting that additional post-entry processes required for productive infection are also affected. Finally, the partial rescue of viral replication by the cell-permeable IP4 analog that was not accompanied by the rescue of the vacuole phenotype (Fig. 3E and 5F) provides direct functional evidence that soluble inositol phosphates themselves contribute to efficient coronavirus infection.

The precise relationship between the antiviral activity of UNC7844, inhibition of the soluble inositol phosphate pathway and phosphoinositide remodeling remains unclear. One possibility is that decreased levels of soluble IPs release the inhibition of various phosphatases as demonstrated for the case of IP7-mediated inhibition of OCRL ^21^. Alternatively, since IPMK itself has a lipid PI3K activity, its direct inhibition by our inhibitors may contribute to the observed lipid remodeling ^22^.

The identity of the soluble inositol phosphate species that are required for coronavirus replication remains to be determined. Although UNC7844 markedly reduced cellular IP5 and IP7 levels (Fig. 2H), the rescue experiments implicate soluble inositol phosphate signaling more broadly, and additional higher-order inositol phosphates may also contribute. Notably, unlike its well-documented role in retroviral capsid stabilization ^3–6^, IP6 is unlikely to mediate the early pre-replication effects of UNC7844 on the coronavirus life cycle, as the drug did not alter IP6 levels (Fig. 2H). The host processes regulated by soluble inositol phosphates also remain incompletely defined. In addition to controlling endosomal trafficking, soluble inositol phosphates may regulate other events required for productive infection, including those associated with viral replication following genome release. Defining these mechanisms will be an important goal for future studies. Irrespective of the precise molecular mechanism, our findings expand the known role of soluble inositol phosphates and their kinases beyond their function in retroviral capsid stabilization and highlight their importance at early stages of coronaviral infection.

From a therapeutic perspective, targeting host pathways represents a complementary approach to direct-acting antivirals, offering a reduced risk of viral resistance and the potential for broad-spectrum activity. This strategy could provide a broadly applicable and rapidly deployable therapeutic approach for future viral outbreaks and pandemics. Our findings establish the soluble inositol phosphate pathway as a druggable host vulnerability and demonstrate that simultaneous inhibition of multiple nodes within this pathway produces substantially greater antiviral efficacy than inhibition of individual kinases. Although additional optimization of UNC7844 will be required to improve its pharmacological properties, our work provides proof of concept that pharmacological targeting of the soluble inositol phosphate network represents a promising strategy for host-directed antiviral therapy.

## Methods

### Cells, viruses and chemicals

The murine 17-CL1 cells (derived from 3T3 cells, BEI Resources, Cat# NR-53719) and LN-18 human glioblastoma cells (ATCC, CRL-2610) were maintained in Dulbecco Modified Eagle Medium (DMEM, Life Technologies, 11965092) supplemented with 10% fetal bovine serum (FBS, Cytiva SH30910.33), and 1% penicillin and streptomycin (Gibco, 15140122). Bone marrow-derived macrophages (BMDMs) were differentiated from bone marrow isolated from femurs of 8-week-old female mice and grown in DMEM supplemented with 10% FBS, 20% L929 conditioned medium and 1% penicillin and streptomycin. THP1 human monocytic leukemia cells (ATCC, TIB-202) were grown in RPMI supplemented with 10% FBS, 50 mM beta-mercaptoethanol and 1% penicillin and streptomycin. Before infections, THP1 cells were differentiated to macrophages (TDMs) using 10 nM PMA for 3 days or 50 nM PMA for 24 hours as indicated on the figure legends. All cells were incubated at 37°C and 5% CO2.

Unless indicated otherwise, all MHV infections in-vitro and in-vivo were with MHV-A59 strain (ATCC, Cat. # VR-764). The recombinant Murine Coronavirus MHV-A59 with enhanced green fluorescent protein (eGFP) (MHV-GFP) was from BEI Resources (NR-53716). MHV-S (ATCC, Cat. # VR-766) was used for the infection of BMDMs. HCoV 229E (ATCC, Cat. # VR-740) was used for the infection of human cells.

GNF-362 was from Cayman Chemicals (35866). The following compounds were from Selleckchem: PI-103 (S1038), PIK-93 (S1489), PI4KIIIbeta-IN-10 (E2981), GDC0980 (S2696), apilimod (S6414).

### Genetic manipulations of the inositol phosphate pathway

For IPMK genetic deletion, CRISPR-Cas9-mediated genetic deletion of all copies of human *IPMK* in human LN-18 cells (ATCC CRL-2610) was performed as previously described ^6, 23, 24^. Briefly, guide RNAs were optimized and selected by the computational tool “CRISPOR” ^25^ to enhance target selectivity for *IPMK* exons 1 and 6 (see supplementary materials for gRNA locations at the *IPMK* locus). IPMK guide RNA sequences for Exon 1 were 5’-CACCGGCGATCGAGTCCACCCCTGA-3’ (A, Forward) and 5’-AAACTCAGGGGTGGACTCGATCGCC-3’ (B, Reverse). Exon 6 guide RNAs were 5’-CACCGCCAAGATGTATGCGCGTCAC-3” (C, Forward) and 5’-AAACGTGACGCGCATACATCTTGGC-3’ (D, Reverse). Oligonucleotide pairs “A/B” and “C/D” were homologous to portions of *IPMK* exon 1 and 6, respectively, these pairs were annealed and cloned into the BbsI site of pX459v2 ^26^, resulting pX459v2-*IPMK*-exon1 and pX459v2-*IPMK*-exon6, which were co-transfected into human LN-18 cells (ATCC CRL-2610). Transfectants were selected for using 1 µg/mL puromycin for two days followed by selection with 0.75 µg/mL puromycin for one day, antibiotic was then removed followed by single cell cloning in 96-well plates. Wells containing one colony were expanded and screened for IPMK knockout (IKO) by western blot. IKO was confirmed by analytical PCR using genomic extracted DNA as shown in supplementary data (Fig. S1) and the sequence of all alleles was determined by Sanger sequencing.

For the expression of wild type or kinase-dead IPMK, N-terminally HA-tagged IPMK (WT or the kinase deficient D144A mutant were cloned into pLPCX (Clontech). VSVg pseudotyped MLV was prepared by transfecting pLPCX-HA-IPMK along with pMLV-Gag-Pol and pVSVg into HEK293T cells using PolyJet transfection reagent (SignaGen). Released virus was harvested and HA-IPMK wild type (WT) or D144A viruses were used to infect IPMK-knockout (IKO) human LN-18 cells (clone 2-G9) at MOI of less than 1 as described previously ^23, 24^. Control cells were transduced with empty vector, selection for puromycin resistance was initiated at 3 days post infection after which cells were maintained in puromycin-containing medium. Cells were tested by western blot for expression of the IPMK in whole cell extracts.

For IPMK knockdown, 17-Cl1 cells were reverse-transfected with ON-TARGETplus Mouse Ipmk siRNA SMARTPool (Horizon, L-062885-00-0005) using Lipofectamine RNAiMAX (ThermoFisher Scientific, 13778075). Infection with MHV-A59 was performed 48 hours later.

For IP4-PM treatments, 17-Cl1 cells were infected with MHV-GFP at an MOI of 2 for one hour followed by virus removal, wash and treatment with 10 μM UNC7844 or DMSO. At 3 hpi, medium was replaced to 10 μM IP4-PM (SiChem, 4-2-1345) or its vehicle control (0.01% Pluronic F-127 and 0.1% DMSO) for 1.5 hours. Then, medium was replaced to fresh 10 μM UNC7844 or DMSO control. RNA was collected at 7 hpi.

### Protein Expression and Purification

#### IP3K, IPMK and IP6K Kinase Activity Assay

Recombinant human IPMK, IP3KA, and IP6K2 were prepared as previously described ^12^. For each assay, enzyme concentrations were optimized by serial dilution to ensure measurements were conducted within the linear range of activity.

For the IP3K kinase activity assay, each reaction contained 1.0 μM 1,4,5-InsP3 and [³H]-1,4,5-IP3 (approximately 30,000 dpm; American Radiolabeled Chemicals, Inc., ART-0270) in a 100 μL incubation buffer comprising 20 mM HEPES (pH 7.2), 100 mM KCl, 3.5 mM MgCl₂, 20 μM EDTA, and 1.0 mM ATP, along with test compound in DMSO (or vehicle control) and 0.25 nM IP3K-A.

For IPMK assays, [³³P]-1,3,4,5-IP₄ was first generated by incubating 1,4,5-IP₃ with [³³P]-ATP and IPMK. The product was isolated using a 30 kDa molecular weight cutoff filter. Each reaction then contained trace amounts (∼400,000 dpm) of [³³P]-1,3,4,5-IP₄ in a 100 μL incubation buffer consisting of 20 mM HEPES (pH 7.2), 100 mM KCl, 3.5 mM MgCl₂, 20 μM EDTA, 1.0 mM ATP, and 1.0 μM 1,3,4,5-IP₄, along with test compound in DMSO (or vehicle control) and 2 nM IPMK.

For the IP6K kinase activity assay, each reaction contained 1.0 μM IP6 and [³H]-IP6 (approximately 30,000 dpm; American Radiolabeled Chemicals, Inc., ART-1915) in a 100 μL incubation buffer comprising 20 mM HEPES (pH 7.2), 100 mM KCl, 3.5 mM MgCl₂, 20 μM EDTA, and 1.0 mM ATP, along with test compound in DMSO (or vehicle control) and 3.5 nM IP6K2.

Reactions were quenched after 60 minutes by addition of two volumes of 0.2 M NH₄H₂PO₄ (pH 3.9) containing 20 mM EDTA and stored at 22 °C. Samples were analyzed by HPLC using a Partisphere SAX 120 column (5 μm, 4.6 × 125 mm) with a 250 μL injection volume. Elution was performed using a gradient formed by mixing Buffer A (1 mM Na₂EDTA) and Buffer B (Buffer A supplemented with 2.5 M NH₄H₂PO₄, pH 3.9). Radioactivity was monitored using a β-RAM Model 6 in-line scintillation detector. The eluent flow rate (1.0 mL/min) was mixed with monoflow scintillation fluid at 1.0 mL/min for ³³P or 2.5 mL/min for ³H detection.

Radiochromatography data were collected and analyzed using Laura software (v6.1.2.36). IC₅₀ values were determined using GraphPad Prism (version 11.0.0) by pooling data from at least three independent experiments and are reported with standard deviations.

### Cell Culture and Assay of Intracellular Inositol Phosphates

For inositol phosphate analysis, 1 × 10 17-CL1 cells were seeded in 10 cm dishes and cultured for 3 days in 7 mL of medium containing 10 μCi/mL [³H]inositol (American Radiolabeled Chemicals), reaching approximately 70% confluence. Cells were then incubated with 10 μM inhibitors or vehicle control for 18 h.

Reactions were quenched by acidification, and inositol phosphates were extracted and separated by HPLC using a Synchropak Q100 column (250 mm × 4.6 mm), as described above. Fractions were collected at 1-minute intervals over 66 minutes, mixed with 2.5 mL Monoflow-4 scintillation fluid (National Diagnostics), and radioactivity was quantified using a scintillation counter.

### X-ray Crystallography Structural Studies

Crystals of human apo-IPMK were prepared as previously described ^27^. Ligand-bound complex crystals were generated by soaking apo crystals in a solution containing 2–10 mM compound, 35% (w/v) PEG 400, 0.1 M Li₂SO₄, and 100 mM HEPES (pH 7.5) at 25 °C for 3 days.

Diffraction data were collected at APS beamlines 22-ID and 22-BM and processed using HKL2000. Structures were determined by rigid-body refinement followed by difference Fourier synthesis and refined using programs within the CCP4 suite. Model quality, including rotamer outliers and clash scores, was assessed using Phenix. Molecular graphics were prepared with PyMOL (Schrödinger, LLC).

Atomic coordinates and structure factors for the human IPMK–compound co-complexes have been deposited in the Protein Data Bank under accession codes 35WJ (UNC7467) and 35WK (UNC7844). Detailed data collection and refinement statistics are provided in Supplementary Table 1.

AlphaFold-predicted full-length structures of IP3KA and IP6K2 were manually modeled based on the UNC7844-bound IPMK structure to reflect potential ligand binding.

### Synthesis of UNC7844

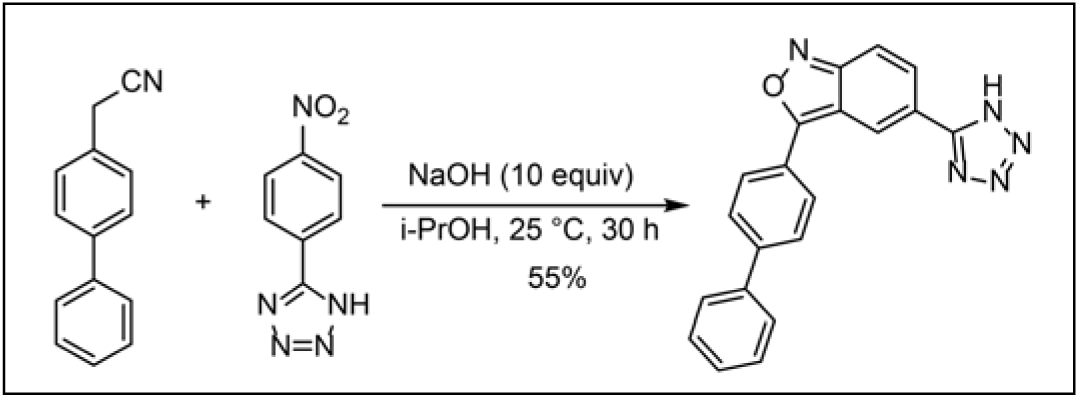

3-([1,1’-Biphenyl]-4-yl)-5-(1H-tetrazol-5-yl)benzo[c]isoxazole (UNC7844)

NaOH pellets were granulated (80.0 mg, 2.00 mmol) and added in i-PrOH (1.0 mL). The mixture was ultrasonicated until NaOH was totally suspended. 5-(4-Nitrophenyl)-1*H*-tetrazole (38.2 mg, 0.200 mmol) was added and the reaction mixture was kept stirring until the solid was dissolved. [1,1’-Biphenyl]-4-yl)acetonitrile (77.3 mg, 0.400 mmol) was added to the reaction mixture. The reaction was stirred at room temperature for 30 h and then acidified by HOAc to pH 4. A light-yellow solid was precipitated and collected by filtration. The filter cake was washed with small amount of EtOAc. The crude product was further purified by reversed ISCO chromatography to afford the title compound (37.4 mg, 0.110 mmol, 55%) as a yellow solid. 1H NMR (400 MHz, DMSO-d6) δ 8.81 (d, J = 1.3 Hz, 1H), 8.32-8.18 (m, 2H), 8.07 (dd, J = 9.3, 1.4 Hz, 1H), 7.99 (d, J = 8.4 Hz, 2H), 7.91 (dd, J = 9.5, 1.0 Hz, 1H), 7.84-7.76 (m, 2H), 7.54 (t, J = 7.5 Hz, 2H), 7.50-7.41 (m, 1H). 13C NMR (101 MHz, DMSO-d6) δ 165.74, 157.01, 155.04, 142.74, 138.75, 129.64, 129.18, 128.42, 127.85, 127.30, 126.90, 125.83, 121.05, 120.99, 116.52, 113.67. MS (ESI) for [M+H]+ (C20H14N5O+): calcd. m/z 340.12; found m/z 340.10; LC-MS: 98% purity.

### Snapshot pharmacokinetic (PK) study

PK study was performed at Pharmaron. All the procedures related to animal handling, care, and the treatment in this study were performed according to guidelines approved by the Institutional Animal Care and Use Committee (IACUC) of Pharmaron (PK-R-06012023 and PK-M-07182023) following the guidance of the Association for Assessment and Accreditation of Laboratory Animal Care (AAALAC).

Groups of 2 male CD1 mice were dosed intravenously or orally with 3 mg/kg UNC7844 in 10% NMP, 5% Solutol, 30% PEG-400 in normal saline (v/v/v/v, 10:5:30:55). From each mouse, blood samples (30 µL/sample) were collected from orbital vein such that samples were obtained at 0.5, 1, 2, 4 hours post dosing. At each time point blood samples were collected from two mice in labeled micro centrifuge tube containing EDTA-K_2_ as anticoagulant. Plasma samples were separated by centrifugation of whole blood and stored at -75±15°C until bioanalysis. All samples were processed for analysis by precipitation using acetonitrile (ACN) and analyzed with fit for purpose LC/MS/MS method (Below quantifiable limit (BLOQ) was 5.0 ng/mL). Pharmacokinetic parameters were calculated by non-compartmental analysis model using WinNonlin 8.3.

### Coronavirus infections of cells in culture

Cells were seeded at 10^5^ per well of 24-well plate for RNA analysis or 10^4^ per well of 96-well plate for viability assays. Unless indicated otherwise, cells were infected and inhibitor treatments were performed in serum-free DMEM with 1% penicillin and streptomycin.

### Cell viability assay

Cell viability was determined by CellTiter-Glo assay (Promega) according to manufacturer’s instructions.

### In vivo infections with MHV-A59 and SARS-CoV-2 and treatments with UNC7844

MHV-A59 infections were approved by the Institutional Animal Care and Use Committee of the National Institute of Environmental Health Sciences. Mice were housed in micro-isolator static cages and were maintained under stringent Specific Pathogen-Free (SPF) conditions, in a controlled environment with constant temperature, humidity, and 12-hour light/dark cycle. They were provided plant-based chow diet (NIH-31, Harlan Laboratories, Madison, WI). The experimental mice were acclimatized for a week before being randomly assigned to experimental groups. 5-month-old female C57BL/6J mice were infected intraperitoneally (i.p.) with 10^6^ pfu MHV-A59 and treated with 25 mg/kg i.p. UNC7844 or vehicle control twice daily. Mice were sacrificed 48 hours post infection. The formulation consisted of 1 mg/ml UNC7844, 0.4 mg/ml Tris base, 5% Solutol HS 15, 5% NMP (N-Methyl-2-pyrrolidone), and 30% PEG 400 in normal saline. The mixture was sonicated at 37 °C for 1 h and warmed at 40 °C until it turned into a clear solution. The formulations of UNC7844 and its vehicle control were stored at -20 degrees in daily aliquots and thawed before injections.

SARS-CoV-2 infections were performed in BSL-3 facilities at Battelle (Columbus, OH) in accordance with the Guide for the Care and Use of Laboratory Animals. 8-week-old male B6.Cg-Tg(K18-ACE2)2Prlmn/J (Jackson lab, #034860) were single-housed and intranasally infected with 1000 TICD50/mouse of SARS-CoV-2 (USA-WAI/2020, BEI) or PBS control. Mice were treated with 10 mg/kg i.p. UNC7844 or vehicle control twice daily for 6 days and surviving mice were euthanized at day 7 for tissue collection. Animals were weighed daily and monitored twice daily for clinical signs which included hunching, ruffled coat, labored breathing and lethargy. Time of discovery of death or time of euthanization of morbid mice was recorded.

### Time-lapse analysis of endosomal viral trafficking

For labeling of MHV-GFP virions, virus-containing supernatant was labeled with 2.5 mM CM-DiI dye (ThermoFisher, C7000) for 2.5 hours at room temperature. Dye was diluted with PBS, and viruses concentrated on 10 kD Amicon ultracentrifugal filters (Millipore, UFC8010) by centrifugation at 6000g at 4 deg for 20 minutes.

17-Cl1 cells were transiently transfected with GFP-SARA (subcloned into Clontech GFP-C1 from pCMV5B-FLAG-SARA which was a gift from Jeff Wrana (Addgene plasmid # 11738) or GFP-LAMP1 ^28^ plasmids using Lipofectamine 3000 (Thermo Scientific). 48 hours later, cells were replated at 105 per chamber of 8-chamber glass-bottom slide (Thermo Scientific, 155409). Next day, cells were infected with CM-DiI-labeled MHV-GFP. Viruses were removed at 1 hpi, cells were washed once and medium replaced to serum-free medium. At 3 hpi, 10 μM UNC7844 or DMSO control were added and timelapse microscopy was performed on Zeiss LSM 880 Confocal Laser Scanning Microscope at 37°C with 5% CO2 for 3 hours using 40x water objective. Cells with medium GFP intensity were chosen to avoid overexpression artifacts.

### Immunofluorescence staining

For co-staining of nsp2/3 and dsRNA, cells were fixed with 4% formaldehyde for 10 minutes at room temperature, permeabilized with 0.5% Triton X-100 in PBS for 5 minutes and stained with a mixture of anti-nsp2/3 (1:1000, gift from Dr. Susan Baker, Loyola University) and anti-dsRNA (1:100, Millipore, MABE1134) followed by goat anti-rabbit IgG H&L (Alexa Fluor® 488, ThermoFisher) and goat anti-mouse IgG H&L (Alexa Fluor® 594, ThermoFisher).

For PtdI staining, cells were fixed and stained according to Echelon Biosciences protocol for immunocytochemistry (ICC) with lipid antibodies using procedure for Golgi apparatus staining. The following antibodies from Echelon Biosciences were used: anti-PI(4)P (Z-P004, 8 mg/ml); anti-PI(3,4)P2 (Z-P034, 5 mg /ml); anti-PI(3,4,5)P3 (Z-P345b, 10 mg /ml); anti-PI(4,5)P2 (Z-G045, 10 mg /ml); anti-PI(3)P (Z-P003, 5 mg /ml); anti-PI (3,5)P2 (Z-P035, 5 mg /ml) followed by anti-mouse IgG AF594 (1:700, ThermoFisher) or anti-mouse IgM DyLight594 (1:500, ThermoFisher). Nuclei were stained with 5 ug/ml DAPI. Images were acquired with the confocal Zeiss LSM 880 microscope using 20X objective.

### RNA isolation, quantitative real-time PCR and RNA-seq analysis

Total RNA was extracted using the RNeasy kit (Qiagen). For RT-PCR analysis, RNA was reverse transcribed using High-Capacity cDNA Reverse Transcription kit (ThermoFisher Scientific). Quantitative real-time PCR (qPCR) assays were conducted on a CFX96 or CFX Opus 384 real-time PCR instruments (Bio-Rad) utilizing iQ SYBR Green SuperMix (Bio-Rad). Primer sequences can be found in Supplementary Table 6.

For RNA-seq analysis, libraries were prepared using the TruSeq Stranded Total RNA kit (Illumina, San Diego, CA). The indexed samples were sequenced on the Nova-seq 6000 (Illumina) employing a 75-bp single-end protocol as per the manufacturer’s guidelines. Subsequently, reads (ranging from 40 to 80 million reads per sample) were aligned to mm10 reference genome separately using the STAR aligner (version 2.6)^29^. Gene quantification based on GENCODE annotation release (GRCh37, p13) was conducted utilizing Subread featureCounts (version 1.4.6)^30^. Principal Component Analysis (PCA) aided in detecting any potential sample outliers. Comparisons between the vehicle and UC7844-treated infected samples were conducted in DESeq2 to detect differentially expressed genes ^31^. To adjust for the false discovery rate (FDR), the Benjamini and Hochberg method was employed. A gene was deemed significantly differentially expressed if the FDR-adjusted p-value for differential expression was below 0.05.

Hierarchical clustering of the differentially expressed genes was performed in Cluster 3.0^32^ and visualized in JavaTreeview ^33^.

All pathway enrichment analysis were preformed using generated differential gene lists in G:Profiler ^34^.

### Western blotting

Cellular pellets were lysed as described in ^23^. 15 μg total protein were loaded per lane, separated in a 10% NuPAGE Bis-Tris gel (Invitrogen) with 1x MOPS SDS-PAGE buffer (Invitrogen) followed by transfer onto nitrocellulose membranes in 1x NuPAGE transfer buffer with 10% methanol (Invitrogen), the membrane blocked for primary antibodies against IPMK (Rabbit anti-IPMK, 1:5000, gift from Prof. Ashok Venkitaraman) in 5% BSA for 2 hours, or blocked for 1 hour in 5% milk for antibodies against Actin (Rabbit Ab1801, 1:1000, Abcam), both IPMK and Actin antibodies were diluted in their respective blocking buffers and probed overnight, then washed using PBS with 0.1% Tween 20 and detected using HRP-conjugated secondary antibodies and HRP SuperSignal Femto (#34094 Thermo Fisher Scientific) on a ChemiDoc MP (BioRad).

### Lipidomics analysis

Methyl tert-butyl ether (1 mL) and equisplash (1.5 μg/mL in methanol, 0.3 mL) were added to each vial containing approximately two million 17-Cl1 cells at room temperature. Samples were vortexed for 10 min and then transferred to a 2 mL Eppendorf tube. Water (0.2 mL) was added to facilitate phase separation and after a brief vortex, the extracts were centrifuged at 2,000 rcf for 10 min. The top layer was manually removed via pipet, dried to completion, and reconstituted in 100 µL of isopropanol (0.1 mL) for analysis using a Q Exactive HF-X (ThermoFisher, Bremen, Germany) mass spectrometer coupled with a Waters Acquity UPLC liquid chromatograph system. Samples were introduced via a heated electrospray ionization (HESI) source using polarity switching at a flow rate of 0.2 mL/min. Data acquisition and analysis were performed using Xcalibur software. LipidSearch Software (v 5.0 ThermoFisher, Breman, Germany) was used to automatically identify lipids for untargeted lipidomics.

### Statistics and Reproducibility

Values are presented as mean ± standard error of mean (SEM) from a minimum of three independent experiments or biological replicates, unless specified differently in the figure legend. Outliers were excluded from subsequent analyses using the ROUT method ^35^.Two-tailed, unpaired, Student’s t-test was utilized to evaluate significant differences between two means ^36^. Statistical significance was set at p<0.05. Differences between the means with more than two comparison groups were analyzed by either two-tailed unpaired Student’s t test with Welch’s correction or two-way ANOVA with Holm-Sidak correction for multiple comparisons and adjusted p-values were reported. Data analyses were performed using Prism Software 10.0 (GraphPad) or Microsoft Office Excel (Version 16.79.1). No methods were applied to ascertain whether the data met the assumptions of the statistical approach (e.g., normal distribution test).

## Data availability

Lipidomics data are provided in Supplementary Table 4.

Atomic coordinates and structure factors for the human IPMK–compound co-complexes have been deposited in the Protein Data Bank under accession codes 35WJ (UNC7467) and 35WK (UNC7844).

RNA-seq data are available in the Gene Expression Omnibus repository at the National Center for Biotechnology Information (GSE333157).

## Acknowledgements

We would like to thank NIEHS Viral Core for virus production and plaque assays, NIEHS Fluorescence Microscopy and Imaging Center for help with immunofluorescent and time lapse analyses; NIEHS Epigenomics Core for RNA sequencing; NIEHS Integrative bioinformatics Support Group for RNAseq analysis; NIEHS Structural Biology Core for assistance with crystallographic data collection; Dr. Susan Baker (Loyola University) for anti-nsp2/3 antibody. This research was supported by the Intramural Research Program of National Institute of Environmental Health Sciences of the National Institutes of Health Z01 ES102205 (to X. L.), the University of North Carolina Cancer Research Fund (to X.W), CA258993 and CA293085 (to Q. Z.), NIGMS R35 GM156389 (to R.D.B), and NIEHS contract HHSN273201700005C. The lipidomics core facility at UNC Chapel Hill is partly supported by the National Science Foundation (CHE-1726291).

The contributions of the NIH authors (I. S., H. W., C. G., C. M. G., S. S., R. S., and X. L.) were made as part of their official duties as NIH federal employees, are in compliance with agency policy requirements, and are considered Works of the United States Government. They are not subject to copyright protection within the United States. However, the findings and conclusions presented in this paper are those of the authors and do not necessarily reflect the views of the NIH or the U.S. Department of Health and Human Services.

The contributions of Y. Z., A. C., Q. Z., R. D. B., and X. W. are licensed under the Creative Commons Attribution 4.0 International-NonCommercial (CC BY 4.0 NC) license selected for this preprint.

## Author contributions

I. S. designed and coordinated the study, designed and performed experiments, analyzed data and wrote the manuscript; H. W. designed the study, evaluated UNC7437, UNC7467, UNC7844, performed structural and biochemical experiments, analyzed data, and wrote the manuscript; Y. Z. and X. W. synthesized UNC7437, UNC7467, UNC7844; C. G. performed inositol profiling experiments; A. C. and Q. Z. performed and analyzed lipidomic data; C. M. G. provided GFP constructs for early and late endosomal vesicles and analyzed data; S. S. and R. S. coordinated the study and analyzed data; R. D. B. generated IPMK WT and KO human LN-18 cells as well as stable LN-18 cell lines re-expressing WT or D144A mutant IPMK, coordinated the study, and analyzed data; X. L. guided, designed, and coordinated the study, analyzed data, and wrote the manuscript. All authors critically reviewed the manuscript.

