## Supplementary Figures for "Host soluble inositol phosphate signaling promotes coronavirus replication"

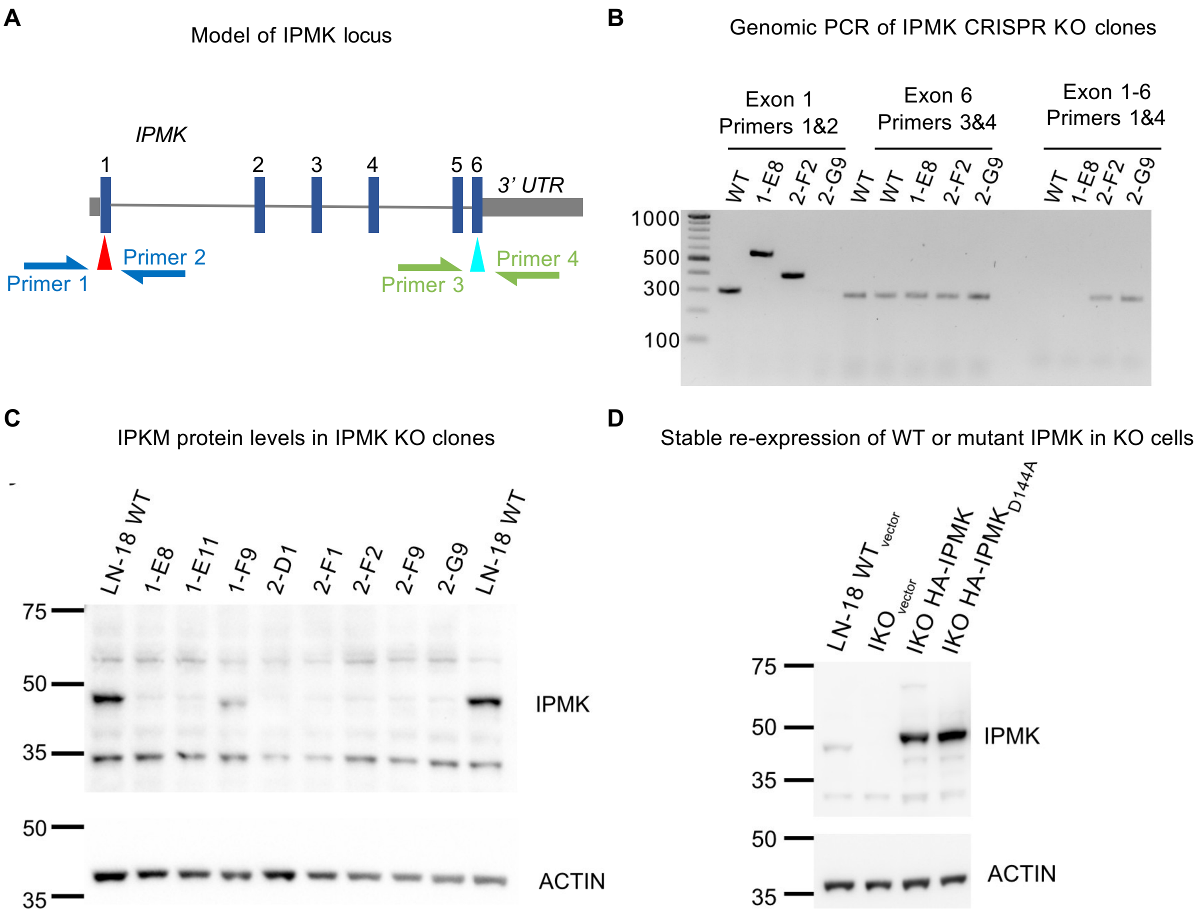


**Figure S1. Creation of IPMK knockout clones and stable re-expression of IMPK in LN-18 cells.**

(A) Model of human *IPMK* genetic locus, showing the intron/exon structure of the gene. Red and teal arrows indicate positions targeted by guide RNAs, which after break repair should produce an Exon 1-6 fusion allele that can be detected in genomic DNA using a 5' primer at Exon 1 and a 3' primer at Exon 6. (B) Analytical PCR of indicated monoclonal human LN-18 cell CRISPR-Cas9 clones tested for IPMK-knockout (KO). PCR reactions of Exon 1 using primers 1 & 2 (left lanes) show clone 2-G9 has no PCR product, PCR of Exon 6 using Primers 3 & 4 (middle lanes) shows PCR products in all lanes, suggesting the entire *IPMK* locus was present in the genomes of all clones tested, and PCR of "Exon 1-6" using Primers 1 & 4 (right lanes) shows presence the Exon1-6 fusion allele resulting from repair of Cas9-induced DNA breaks to exclude exons 2,3,4 and 5 of *IPMK,* thus clone 2-G9 is a validated genetic knockout of *IPMK*. Since these PCR reactions worked as expected, clones containing loss of the *IPMK* exon 1 locus with the presence of a PCR product corresponding to a fusion allele(s) or clear insertion/deletion (in/del) repair events in exon 1 were chosen for western analysis. (C) Western blots of whole cell lysates probed using antibodies directed against endogenous IPMK (upper) or actin (lower), suggesting indicated monoclonal CRISPR cell lines contain no detectable full-length IPMK protein compared to wild-type LN-18 cells, except clone 1-F9. Monoclonal LN-18 cell line 2-G9 was the clone used for complementation with wild-type and kinase-dead IPMK. (D) Western blots of monoclonal LN-18 cell line 2-G9 used for stable complementation with HA-tagged wild-type or kinase-dead (D144A) IPMK. Whole cell lysates were probed using antibodies directed against endogenous IPMK (upper) or actin (lower), showing re-expression of wild-type or kinase-dead IPMK.

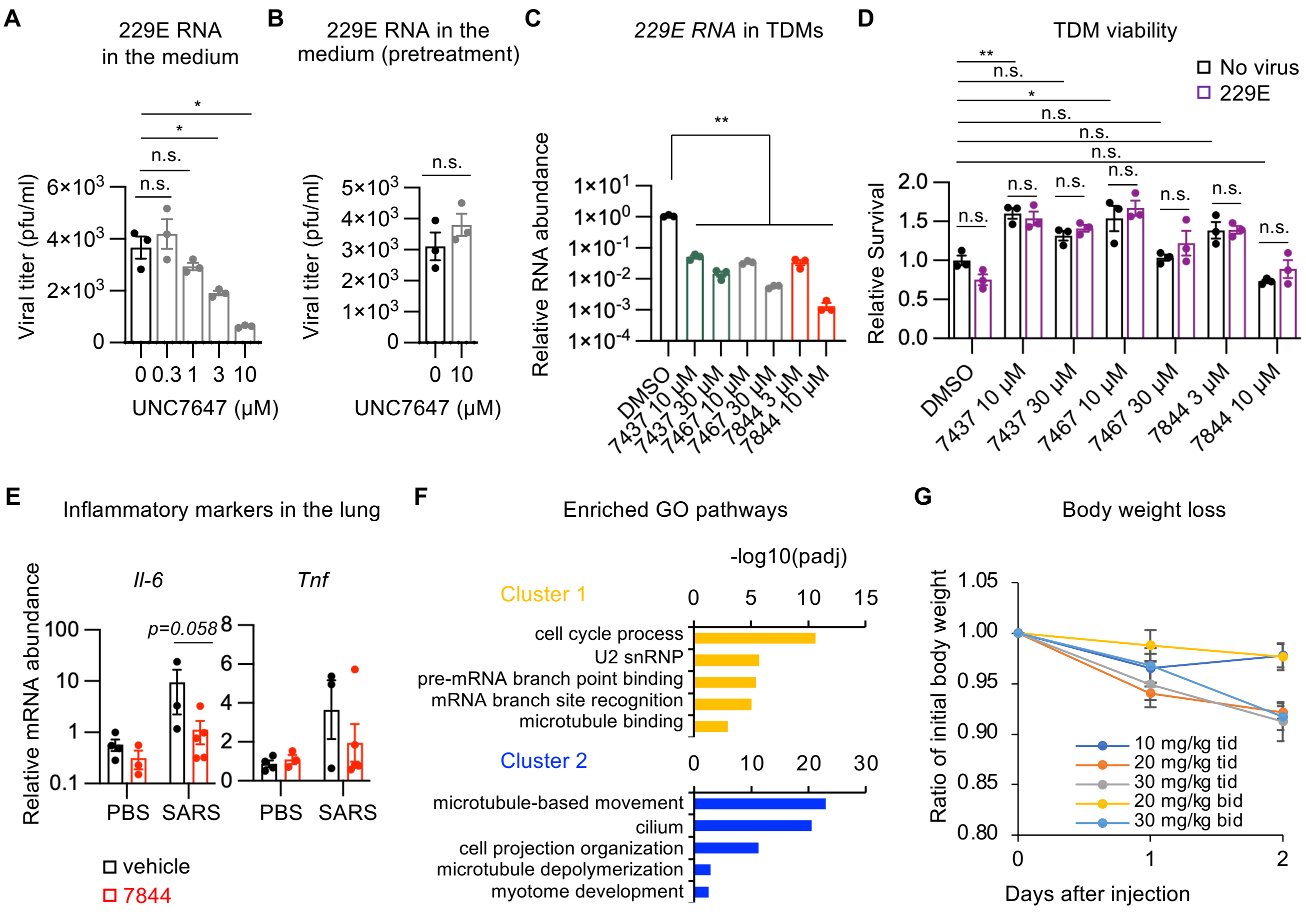

**Figure S2. IPK inhibitors suppress coronavirus replication.**

(A) Dose-response curve of antiviral activity of UNC7467 in THP1 cells. THP1 cells were differentiated with 10 nM PMA for 3 days and then infected with 229E at an MOI of 2. Virus was removed at 2.5 hpi, cells were washed and incubated with the indicated concentrations of UNC7467 for 48 hours in full growth medium containing 10% FBS. Virus in the medium was quantified by qPCR and compared with a stock of 229E with a known pfu (n=3 replicates/group, values are expressed as mean ±SEM, 2-tailed unpaired t test with Welch's correction, *p<0.05, n.s., not significant). (B) UNC7467 does not inhibit 229E replication when used as a pre-treatment. Experiment as in (A) but cells were treated with 10 μM UNC7844 during 3 days of PMA-mediated differentiation before the viral infection (n=3 replicates, values are expressed as mean ±SEM, 2-tailed unpaired t test, *p<0.05, n.s., not significant). (C) THP1 cells were differentiated with 50 nM phorbol myristate acetate (PMA) for 24 hours into TDMs and then infected with 229E at an MOI of 1. Virus was removed at 2 hpi and cells were washed and treated with the indicated inhibitors for 17 hours (n=3 replicates/group, values are presented as mean±SEM, 2-tailed unpaired t test with Welch's correction, **p<0.01). (D) Cell viability of TDMs treated as in (C) was determined by CellTiter-Glo assay at 44 hpi (n=3 replicates/group, values are expressed as mean ±SEM, 2-way ANOVA, *p<0.05, **p<0.01, n.s., not significant). (E-F) UNC7844 shows a trend toward anti-SARS-CoV-2 activity in vivo. hACE2-overexpressing mice were intranasally infected with SARS-CoV-2 and treated with 10 mg/kg twice daily UNC7844 or vehicle control for 7 days. (E) mRNA levels of indicated genes in the lung were analyzed by qPCR (n= 4, 3, 3, and 5 mice, values are expressed as mean ±SEM, 2-tailed unpaired t test between 7844 vs vehicle treated SARS infected mice). (F) Pathway enrichment analysis for the clusters indicated in Figure 3E. (G) Dose escalation of UNC7844 in C57BL/6J male mice. UNC7844 was i.p. injected at indicated doses either three time daily (tid) or twice daily (bid) (n=4 mice/group, values are presented as mean ± SEM).

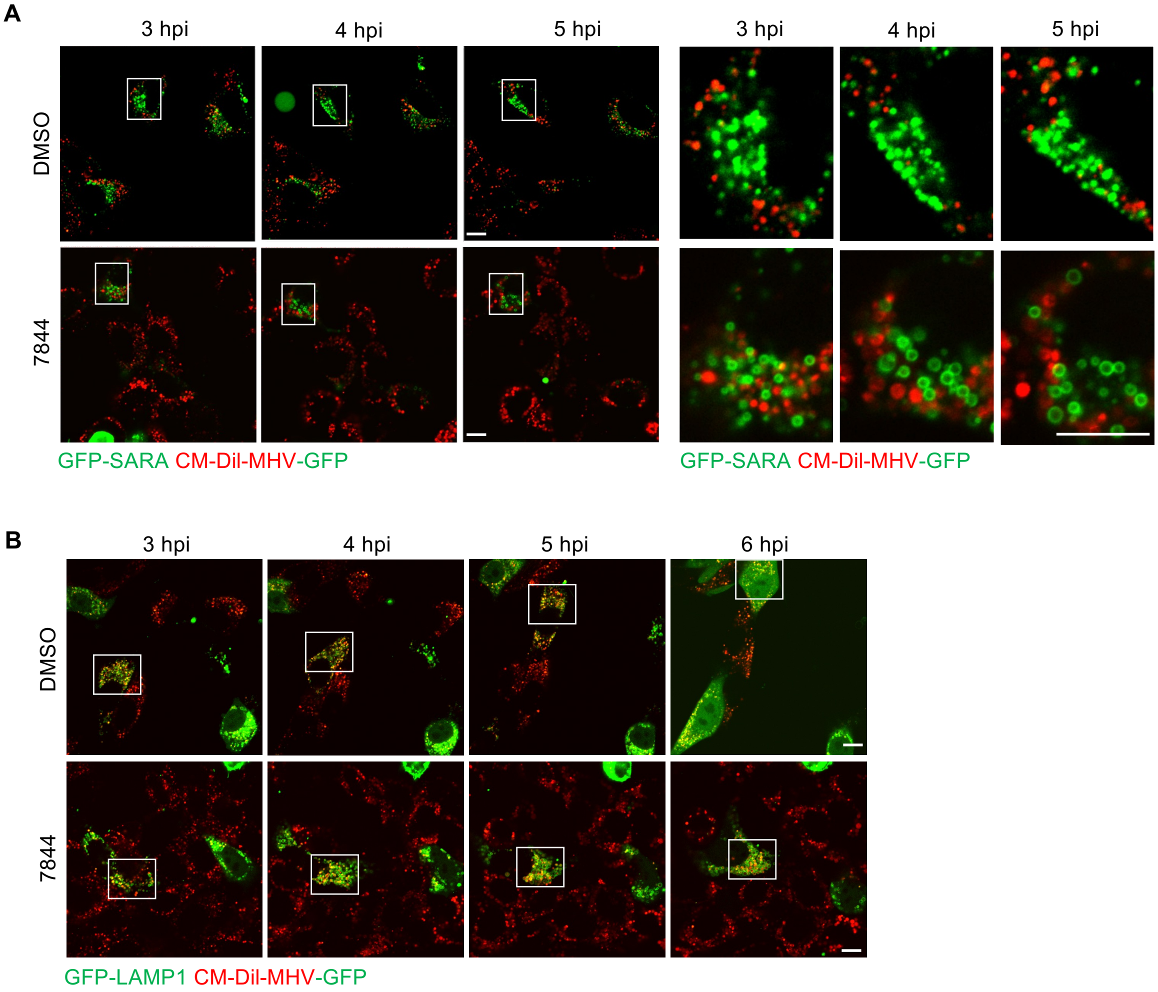


**Figure S3. UNC7844 inhibits endolysosomal viral trafficking.**

(A) Mouse 17-Cl1 fibroblasts transiently transfected with GFP-SARA (early endosome marker) were infected with CM-DiI-labeled MHV-GFP virus for 1 hr, then virus was washed away. At 3 hpi, 10 µM UNC7844 or DMSO were added and cell were analyzed by time lapse microscopy. Right panels, zoom-in of the boxed regions Bars, 10 µm. (B) Mouse 17-Cl1 fibroblasts transiently transfected with GFP-LAMP1 (late endosome/lysosome marker) were treated as in (A). The boxed regions were zoomed-in in Figure 4E. Bars, 10 µm.

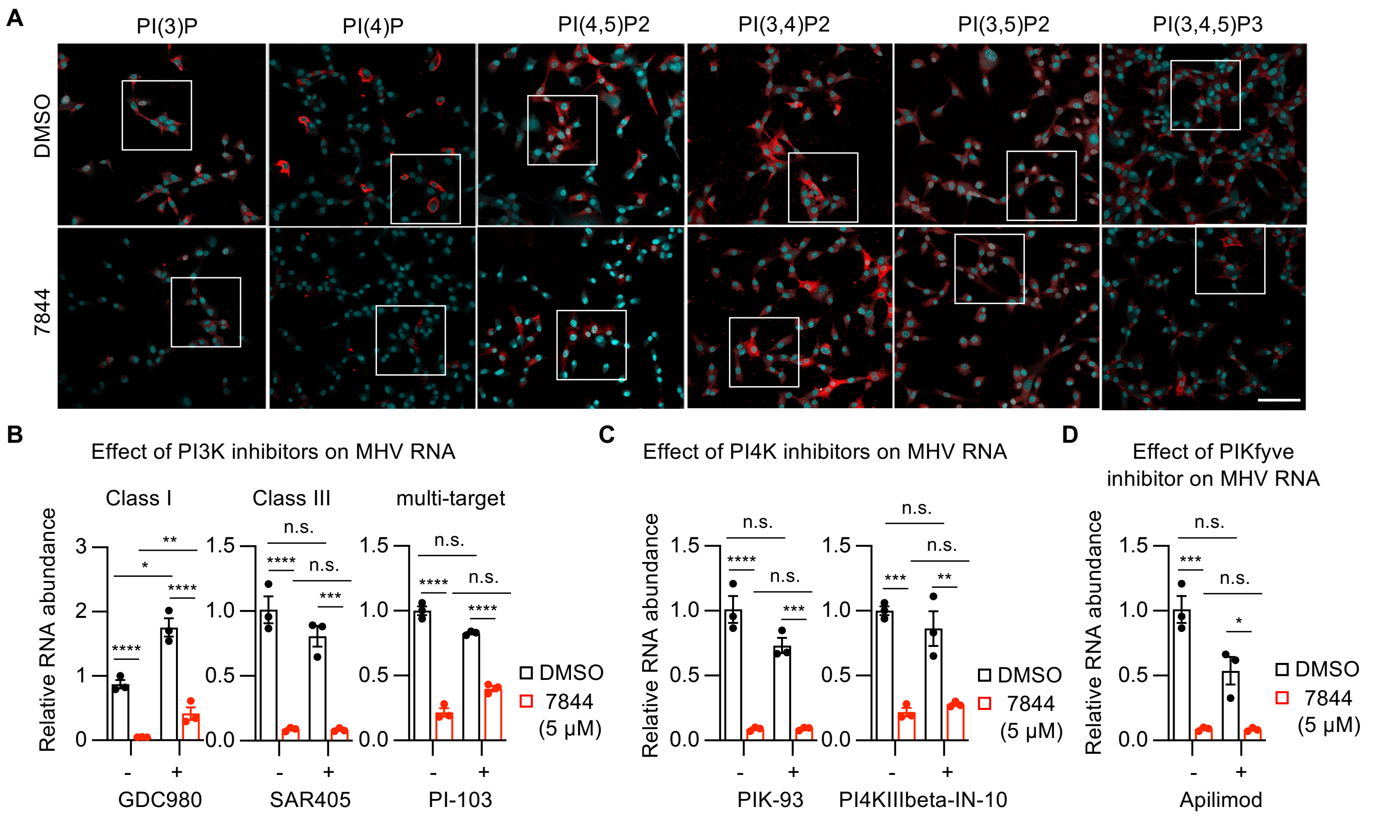


**Figure S4. UNC7844 remodels phosphatidylinositide composition and exhibits substantially greater antiviral potency than selective phosphoinositide kinase inhibitors.**

(A) Immunostaining for specific phosphoinositides. The correspondent boxed regions were zoomed-in in Figure 5C. Bar, 100 μm. (B) 17-Cl1 cells were infected with MHV at an MOI of 3. Virus was removed at 1 hpi, cells were washed and treated with 1 μM SAR405, 100 nM PI-103, 200 nM GDC980 with or without 5 μM UNC7844. Cellular RNA was collected at 7 hpi and MHV RNA was quantified by qRT-PCR (n=3 replicates/group, values are expressed as mean ±SEM, 2-way ANOVA, *p<0.05, **p<0.01, ***p<0.001, ****p<0.0001, n.s., not significant). (C) Experiments as in (B) but using 2 μM PIK93 or 2 μM PI4IIIbeta-IN-10 and RNA collection at 6 hpi (n=3 replicates/group, values are expressed as mean ±SEM, 2-way ANOVA, **p<0.01, ***p<0.001, ****p<0.0001, n.s., not significant). (D) Experiment as in (C) but using 200 nM apilimod (n=3 replicates/group, values are expressed as mean ±SEM, 2-way ANOVA, *p<0.05, ***p<0.001, n.s., not significant).
